# Mapping polarization sensitive optical coherence tomography and ultra-high-field diffusion MRI in the macaque brain

**DOI:** 10.1101/2024.04.19.590326

**Authors:** Pramod K Pisharady, Marilyn Yeatts, Hamza Farooq, Matthew Johnson, Noam Harel, Steen Moeller, Jan Zimmermann, Essa Yacoub, Kamil Ugurbil, Sarah R. Heilbronner, Christophe Lenglet, Taner Akkin

## Abstract

This paper provides comparisons between microstructure and two-dimensional fiber orientations measured optically using polarization sensitive optical coherence tomography (PS-OCT) and those estimated from ultra-high-field diffusion MRI (dMRI) at 10.5T in the macaque brain. The PS-OCT imaging is done at an in-plane resolution of ∼10 microns in and around the thalamus. Whole brain dMRI is acquired at an isotropic resolution of 0.75 mm. We provide comparisons between cross-polarization and optical orientation from PS-OCT with the fractional anisotropy and two-dimensional orientations extracted from dMRI using a diffusion tensor model. The orientations from PS-OCT are also extracted computationally using a structure tensor. Additionally, we demonstrate the utility of mesoscale, PS-OCT imaging in improving the MRI resolution by learning the mapping between these contrasts using a super-resolution Generative Adversarial Network.

## 1 Introduction

Optical coherence tomography (OCT), which produces depth-resolved images at micrometer resolution, has been used to visualize cortical layers and neurons [1-3]. Reflectivity is the conventional contrast used in OCT which portrays the tissue morphology. The birefringence property of myelinated nerve fibers is used in polarization-sensitive OCT (PS-OCT) [4] to create additional contrasts such as cross-polarization and retardance that better distinguish white matter from gray matter, and to visualize axonal tracts [5].

Diffusion MRI (dMRI) is currently the only imaging modality to map axon bundles and whole brain structural connections in-vivo at macroscale resolutions [6,7]. Despite the submillimeter resolution of dMRI available with state-of-the-art ultra-high-field imaging [8], the imaging resolution needs further refinement to capture the complex neural connections formed by the ∼86 billion neurons present in a healthy human brain, and to detect abnormalities in these neuronal connections. One of the ways to improve the resolution is through post processing by applying a previously learned mapping between the (macroscale) MRI data and (microscale) microscopy data [9].

Microscopic methods such as histological staining and polarized light imaging are traditionally used to validate neuronal parameters like the fiber orientation and axon density estimated from dMRI [10-12]. The factors that limit the use of machine learning for learning the structural mapping between dMRI and microscopy data include the lack of large area of coverage in microscopy and the lack of one-on-one correspondence between MRI and (group of) microscopy pixels. To increase the area of coverage a tissue slicer has been integrated into PS-OCT to make a serial optical coherence scanner (SOCS) [13,14] which is used to validate dMRI and PS-OCT-based in-plane fiber orientation estimations [15].

In this work, we present the acquisition and processing of high-resolution PS-OCT data (∼10 micron lateral; ∼5.5 micron axial) using SOCS, from a large block of tissue (25x25x15 mm^3^) around the left thalamus and adjacent regions of the macaque brain. This is made feasible by imaging each slice (the lateral plane of the block) in tiles (4x4) and subsequently combining them through image stitching in post-processing. Multiple imaging contrasts are calculated from the amplitudes and phases of two PS-OCT channels. This includes two-dimensional (2D) optic axis orientation, which gives the in-plane orientation of the axon fibers. The 2D orientations are also computed with a structure tensor analysis of the cross-polarization contrast. Resemblance of the cross-polarization from PS-OCT and fractional anisotropy (FA) from diffusion tensor imaging (DTI) help registering these two datasets, which allows for comparing the PS-OCT-based orientations and 2D orientations extracted from DTI. Finally, we demonstrate the utility of such high-resolution microscopy data in improving the MRI resolution using a Generative Adversarial Network (GAN). We conclude with a discussion of the next steps including the extension to 3D fiber orientations and designing custom deep network models for such data.

## 2 Methods

### 2.1 In-vivo MRI acquisition and processing

The data acquisition in the study was approved by the Institutional Animal Care and Use Committee (IACUC) at our institution. Whole brain dMRI data of the macaque was acquired using a Siemens MAGNETOM whole body 10.5T MRI scanner. The data were acquired at a resolution of 0.75x0.75x0.75 mm^3^, with two b-values 1000 and 2000 s/mm^2^, and 107 diffusion gradient directions with 8 b0 volumes equally interleaved, using a 2D spin echo EPI acquisition with monopolar diffusion weighting. Echo time (TE) and repetition time (TR) used are 65.8 ms and 6000 ms respectively. Two sets of data were collected with reversed phase encoding directions (head to foot and foot to head).

The data were first denoised with the NORDIC algorithm [16] and corrected for distortions due to eddy currents, susceptibility-induced off-resonance artifacts, and subject motion using FSL software [17,18]. A DTI model was subsequently fit to the data using FSL. A mask corresponding to the block of tissue imaged with PS-OCT was created and the FA and primary fiber orientation (v1) maps were extracted from the slices in this block. The 3D orientations were projected to the 2D imaging plane used in PS-OCT.

### 2.2 Ex-vivo PS-OCT acquisition and processing

Perfused and paraformaldehyde-fixed tissue of the macaque brain was blocked in and around the left thalamus resulting in a tissue sample of 25x25x15 mm^3^. This sample was then mounted to a motorized stage with an agar gel support and imaged with the PS-OCT scanner. The exposed face was imaged from the anterior to posterior direction in the coronal plane using 4x4 tiles, each tile approximately 7x7 mm^2^, with 15% overlap. After imaging the slice, a 100-micron thick tissue was removed from the surface to access deeper regions. This resulted in 156 total slices. We excluded the data from a few slices that had light focusing and partial coverage issues, and used the data from 135 slices for subsequent analysis.

The PS-OCT data were processed to extract co- and cross-polarization channel amplitudes that were used to calculate the reflectivity, *R(z)*, and phase retardance, δ(z), contrasts.

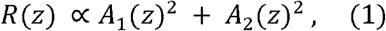

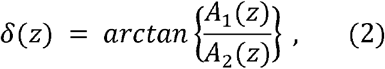

where A denotes the amplitude as a function of depth z and the subscripts represent the polarization channels 1 and 2. Here, *A*_1_ (*z*)^2^ is the cross-polarization contrast. The relative axis orientation, θ′(z), on the other hand, is obtained by using phase (*φ*) information of these PS-OCT channels.

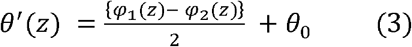

For our polarization-maintaining fiber based system, this measurement has an offset 0_0_which can be removed by utilizing a calibration line [14] or by determining the offset as explained in the next subsection.

For all the contrasts, *en-face* images are calculated with a lateral resolution of 10x10 microns and a slice thickness of 100 microns. A set of 4x4 image tiles per slice is stitched together to one slice using linear blending in MATLAB.

### 2.3 Optical and structure tensor-based orientation estimation

The optical fiber orientations were calculated using Eq.(3). The difference in phase of the two channels provides the relative orientation at each pixel. The offset *θ*_0_ is an arbitrary angle resulting from fluctuations in the system (due to fiber movement, temperature variations etc.) during data acquisition. To obtain *θ*_0_ (which is constant for a slice), local fiber orientations relative to the horizontal axis were estimated using morphological processing of intensity in each en-face slice. A linear structural element with a known length and angle, centered on each pixel, is used for the morphological processing. The angle of this structural element was then varied from 0 to 180 degrees in 5-degree increments. The angle with the maximum number of pixels with an intensity > 0.3 within the length of the structural element was selected as the estimated local fiber orientation. The difference between the estimated orientation and the measured relative optical orientation in a slice forms a Gaussian distribution of offsets. The peak of the distribution was used as the offset *θ*_0_ for the given slice.

The fiber orientations are also calculated with structure tensor analysis [19-21]. The analysis relies on finding the directionality of textures by using the neighboring gradients of pixels. The gradients are calculated using a derivative-of-gaussian filter with a standard deviation of 0.27. The structure tensor of the gradients and its eigenvalues are calculated next. The eigenvalues represent the local gradients of the image intensity and the eigenvectors represent the gradient directions. The eigenvector corresponding to the smallest eigenvalue is taken as the fiber orientation as the fiber axis has the smallest gradient (that is lesser intensity variation along the fiber than across the fiber).

### 2.4 Structural mapping using deep learning

We investigated the potential enhancement of dMRI-derived metrics, particularly FA maps, through the integration of high-resolution PS-OCT data. We used the cross-polarization en-face images as it provided a good contrast that resembles the FA images. We used the PS-OCT imaged block of the brain as the region of interest (ROI) to facilitate direct comparison and analysis. The ROI comprises 27 x 20 pixels/voxels in the dMRI dataset and 2810 x 2740 pixels in the PS-OCT dataset within the corresponding area, with the PS-OCT data possessing 104 x 137 times greater pixel resolution compared to the dMRI data.

With the significantly higher resolution of PS-OCT data, there emerges a promising opportunity to enhance the resolution of FA images. However, the limited dataset comprising only 18 sample MRI slices (corresponding to 135 PS-OCT slices, which are resized to 18 slices using bicubic interpolation in MATLAB) presents a challenge for conventional deep learning methods such as Deep Neural Networks (DNNs) and Convolutional Neural Networks (CNNs), which typically require larger datasets for effective training. To address this limitation, we employed multiple strategies. Firstly, we focused on enhancing FA only by a factor of 8 using down-sampled PS-OCT images. Secondly, we did transfer learning of a pre-trained Enhanced Super-Resolution Generative Adversarial Networks (ESRGAN) [22] model, which was initially designed for a maximum of 4 times super-resolution, to adapt it to achieve 8 times enhancement in FA maps using high-resolution PS-OCT data.

The model architecture is the pre-trained ESRGAN [22] as a base, in a frozen state, and a new super-resolution layer comprising a convolutional layer with 3 filters of 3x3 kernel size. With 3 input channels, this layer incorporated 27 trainable weights per channel along with 3 bias terms per filter, totaling 84 trainable parameters. For optimization, a perceptual loss function computed from features extracted by a pretrained VGG16 [23] convolutional neural network was employed, alongside the Adam optimizer [24] with a learning rate set at 0.1. Training was conducted on 20 epochs with a batch size of 6 high- and low-resolution images, using a 10% validation split for performance monitoring, achieving a minimum training loss of 0.0816. Evaluation included assessing Structural Similarity Index Measure (SSIM) loss, with a final value of -0.0109 indicating a favorable resemblance between generated and the PS-OCT images. The implementation was done using the TensorFlow framework, without data augmentation in training.

## 3 Results

### 3.1 PS-OCT data and its comparison to MRI

Figure 1 shows the en-face cross-polarization images that are calculated by averaging the 3D slices in the axial (z) direction. The results are selected from the sample slices imaged anterior to posterior. In-plane resolution is 10x10 microns and the slice thickness of en-faced images is 100 microns. We used the cross-polarization images for the structure tensor-based orientation estimation and to demonstrate the enhancement of FA images. A movie showing the cross-polarization in all the slices are included in the supplementary material.

**Fig. 1.**
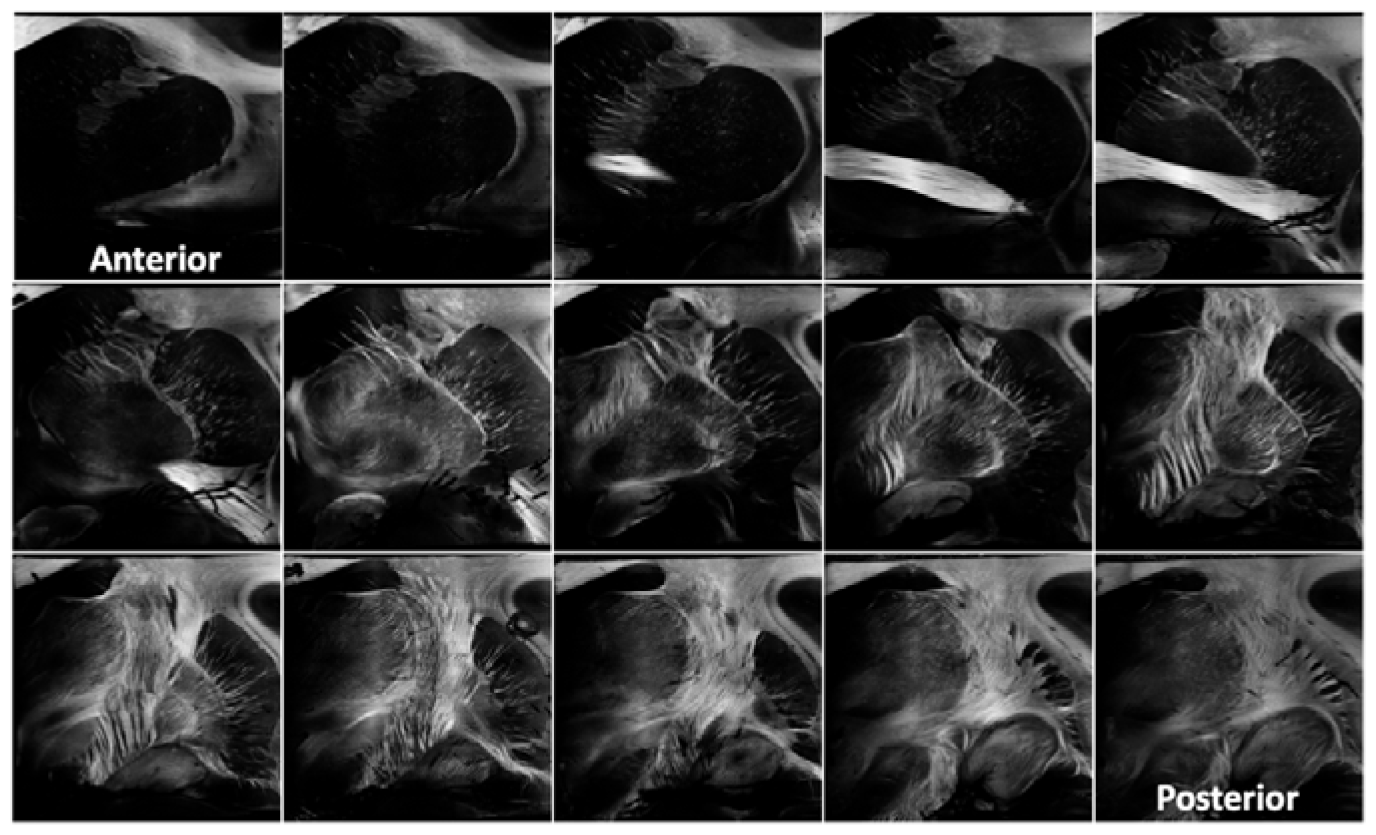
PS-OCT en-face cross-polarization images acquired from a region of interest around the left thalamus of the macaque brain, from anterior (top left) to posterior (bottom right). The order of the slices is left to right and top to bottom.

Figure 2 presents the en-face PS-OCT images including the cross-polarization, retardance, reflectivity and optical orientation, as well as the structure tensor-based orientations that are calculated from the cross-polarization images. The figure also shows a comparison to FA and DTI-based orientation estimates. There is an agreement between the optical orientation and the structure tensor-based orientation, but the optical orientation has higher resolution and appears to be more accurate as it does not depend on imaging features or gradients, which is the case for structure tensor computation. Cross-polarization and retardance contrasts compare well with FA despite the big difference in image resolution (>100x). A second movie showing the optical orientation in all the slices are included in the supplementary material.

**Fig. 2.**
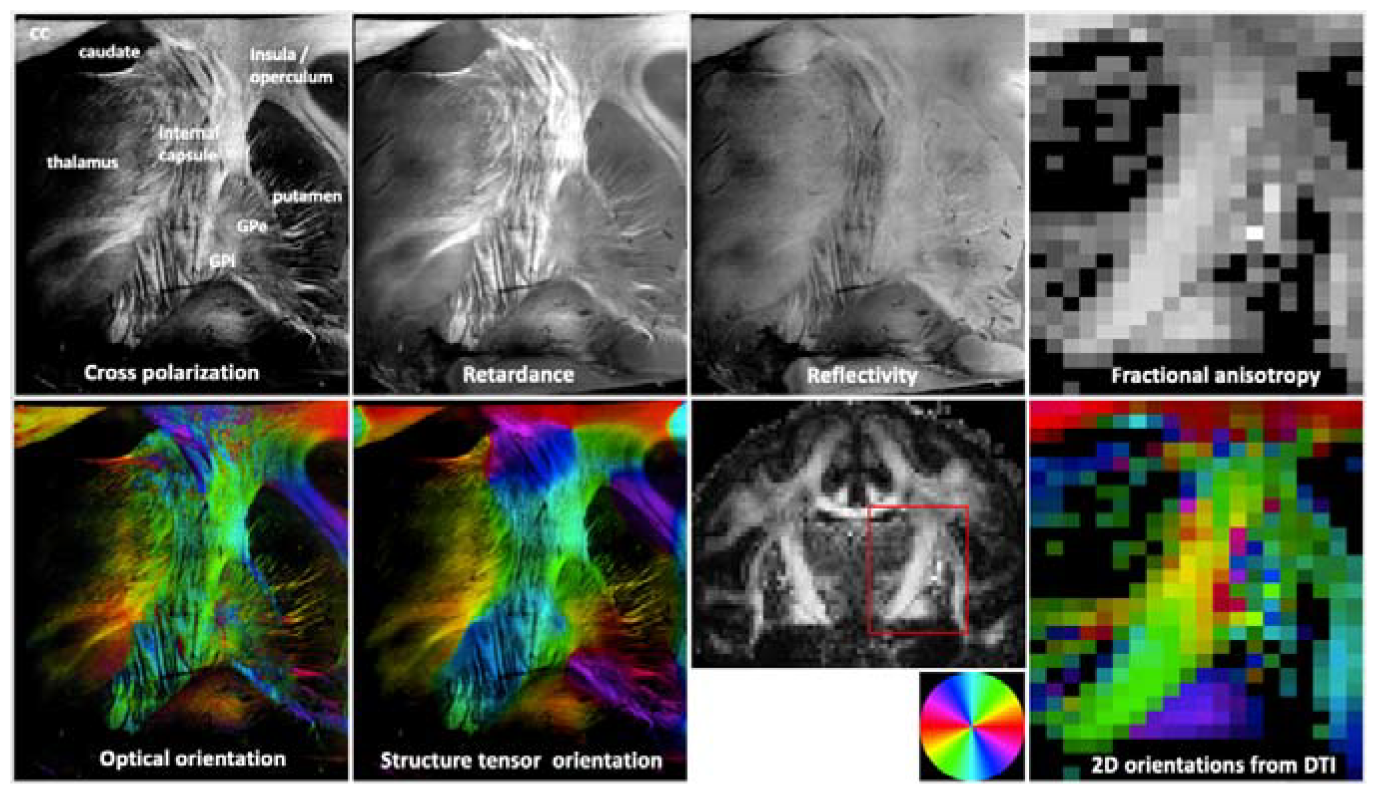
Different imaging contrasts from PS-OCT and their comparison to DTI measures. A coronal view of the FA map highlighting the ROI imaged in PS-OCT and the color coding used for orientation estimates are also shown (second row, third column). CC-corpus callosum, Gpe-globus pallidus externus, Gpi - globus pallidus internus.

### 3.2 Super-resolution DTI by learning mapping between DTI and PS-OCT

Here we demonstrate the effectiveness of Generative Adversarial Network (GAN)- based super-resolution in producing high-quality images from low-resolution DTI data using transfer learning with PS-OCT data (cross-polarization). We used 18 high-resolution PS-OCT images (216 x 160 pixels, down sampled from the full resolution, see the discussion on this in the next section) to achieve an 8 times super-resolution of dMRI data, originally 27 x 20 pixels/voxels.

Figure 3 shows the original FA map, the super-resolution FA map, and its comparison to ground truth PS-OCT (cross-polarization) data. In comparison, the transfer learning of the ESRGAN produced super-resolution FA maps with significant improvement (8x) in the resolution of original FA maps. Enhanced details are visible within the internal capsule, pallidum, putamen, and anterior commissure in the super resolution FA compared to original FA.

**Fig. 3.**
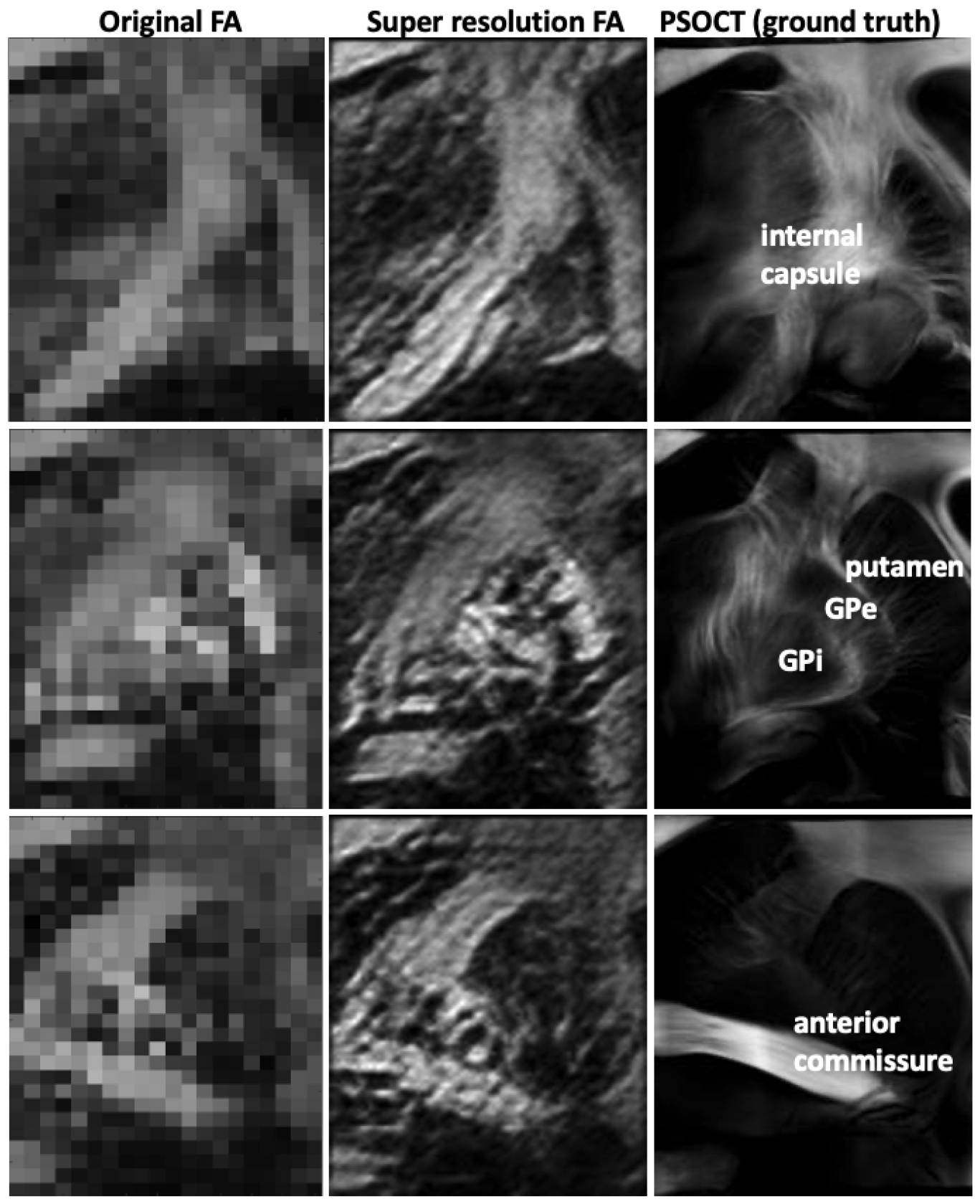
Comparisons between original FA map (left column, 27x20 pixels) and super resolution FA map (middle column, 216x160 pixels) learned through transfer learning of ESRGAN. Ground truth used is the en-face cross-polarization images from PS-OCT (right column, 216x160 pixels). Gpe - globus pallidus externus, Gpi - globus pallidus internus.

## 4 Discussion and Conclusion

We presented preliminary results from the comparison of ex-vivo PS-OCT and invivo dMRI. Both dMRI and PS-OCT data we presented are unique and include high resolution dMRI acquired at 10.5T with 750 microns isotropic resolution. Also, this is the first application of the SOCS technique on a considerable size of macaque midbrain.

Our study aimed to enhance the resolution of dMRI-derived metrics by incorporating information from PSOCT data, rather than attempting to generate PS-OCT images from dMRI. This strategy was motivated by constraints in data size and the substantial resolution disparity between the two imaging modalities (>100x). Therefore, our approach partially leveraged the rich information offered by PS-OCT data to enhance tissue structural details (up to 8x) in FA maps. We accomplished this through transfer learning applied to the pre-trained ESRGAN model. This method enabled us to effectively preserve the anatomical realism in the generated high-resolution FA maps, through learning from PS-OCT data.

In this work we could not leverage the full benefit of microscopic resolution of PS-OCT as we increased the resolution of FA maps only by a factor of 8, by using down sampled version of PS-OCT data. In the future, we plan to enhance the super-resolution factor by exploring customized GANs and Diffusion-based model architectures [25] tailored specifically to our dataset. We plan to develop models capable of achieving super-resolution factors ranging from 10 to 15 times for direct mapping from dMRI resolution. Moreover, we expect that integrating all the diffusion-weighted measurements, which consist of 115 volumes, into the training process will greatly improve model learning. By harnessing the extensive resolution potential offered by PS-OCT data, we aim to generate precise high-resolution diffusion images, thereby pushing forward the capabilities of imaging techniques in neuroimaging research.

Additional data from the other hemisphere of the same macaque is going to be acquired to further these experiments and also to extend the work to fiber tractography from PS-OCT data. We plan to use a calibration line in this data acquisition to facilitate dynamic removal of the offset from the orientation measurement, which will eliminate the need for calculating the orientation offset from morphological signal processing. Our future studies will involve PS-OCT acquisitions from an entire hemisphere or whole macaque brain, to support brain-wide connectome studies at the mesoscale, through hardware and software upgrades. Our future work would also include estimating 3D orientations from PS-OCT data by combining the 2D optical orientations with structure tensor-based third angle (inclination angle) estimation.

## Acknowledgements

This work was supported by NIH grants UM1NS132207, S10 RR029672, P41 EB027061, R01MH126923, and R01MH118257. This work was also supported by CZI grants DAF2020-225709 and DAF2023-329645 from the Chan Zuckerberg Initiative DAF (grant DOI https://doi.org/10.37921/016061nlotjw), an advised fund of Silicon Valley Community Foundation (funder DOI 10.13039/100014989).

## References

1. Srinivasan VJ, Radhakrishnan H, Jiang JY, Barry S, Cable AE. Optical coherence microscopy for deep tissue imaging of the cerebral cortex with intrinsic contrast. Opt Express. 2012 Jan 30;20(3):2220–39.

2. Magnain C, Augustinack JC, Reuter M, Wachinger C, Frosch MP, Ragan T, Akkin T, Wedeen VJ, Boas DA, Fischl B. Blockface histology with optical coherence tomography: a comparison with Nissl staining. Neuroimage. 2014 Jan 1;84:524–33.

3. Magnain C, Augustinack JC, Konukoglu E, Frosch MP, Sakadžić S, Varjabedian A, Garcia N, Wedeen VJ, Boas DA, Fischl B. Optical coherence tomography visualizes neurons in human entorhinal cortex. Neurophotonics. 2015 Feb 9;2(1):015004.

4. de Boer JF, Milner TE, van Gemert MJ, Nelson JS. Two-dimensional birefringence imaging in biological tissue by polarization-sensitive optical coherence tomography. Opt Lett. 1997 Jun 15;22(12):934–6.

5. Wang H, Black AJ, Zhu J, Stigen TW, Al-Qaisi MK, Netoff TI, Abosch A, Akkin T. Reconstructing micrometer-scale fiber pathways in the brain: multi-contrast optical coherence tomography based tractography. Neuroimage. 2011 Oct 15;58(4):984–92.

6. Basser PJ, Pajevic S, Pierpaoli C, Duda J, Aldroubi A. In vivo fiber tractography using DT-MRI data. Magn Reson Med. 2000 Oct;44(4):625–32.

7. Jbabdi S, Sotiropoulos SN, Haber SN, Van Essen DC, Behrens TE. Measuring macroscopic brain connections in vivo. Nat Neurosci. 2015 Nov;18(11):1546–55.

8. Uğurbil K, Xu J, Auerbach EJ, Moeller S, Vu AT, et al., Pushing spatial and temporal resolution for functional and diffusion MRI in the Human Connectome Project. Neuroimage. 2013 Oct 15;80:80–104.

9. Liang Z, Zhang J. Mouse brain MR super-resolution using a deep learning network trained with optical imaging data. Front Radiol. 2023 May 15;3:1155866.

10. Leergaard TB, White NS, de Crespigny A, Bolstad I, D’Arceuil H, Bjaalie JG, Dale AM. Quantitative histological validation of diffusion MRI fiber orientation distributions in the rat brain. PLoS One. 2010 Jan 7;5(1):e8595.

11. Mollink J, Kleinnijenhuis M, Cappellen van Walsum AV, Sotiropoulos SN, et al., Evaluating fibre orientation dispersion in white matter: Comparison of diffusion MRI, histology and polarized light imaging. Neuroimage. 2017 Aug 15;157:561–574.

12. Schilling KG, Janve V, Gao Y, Stepniewska I, Landman BA, Anderson AW. Histological validation of diffusion MRI fiber orientation distributions and dispersion. Neuroimage. 2018 Jan 15;165:200–221.

13. Wang H, Zhu J, and Akkin T. Serial optical coherence scanner for large-scale brain imaging at microscopic resolution. NeuroImage, 84 (2014), 1007–1017.

14. Liu CJ, Williams KE, Orr HT, Akkin T. Visualizing and mapping the cerebellum with serial optical coherence scanner. Neurophotonics. 2017 Jan;4(1):011006.

15. Wang H, Zhu J, Reuter M, Vinke LN, Yendiki A, Boas DA, Fischl B, Akkin T. Crossvalidation of serial optical coherence scanning and diffusion tensor imaging: a study on neural fiber maps in human medulla oblongata. Neuroimage. 2014 Oct 15;100:395–404.

16. Moeller S, Pisharady PK, Ramanna S, Lenglet C, Wu X, Dowdle L, Yacoub E, Ugurbil K, Akçakaya M. NOise reduction with DIstribution Corrected (NORDIC) PCA in dMRI with complex-valued parameter-free locally low-rank processing. Neuroimage. 2021 Feb 1; 226:117539.

17. Andersson JLR, Sotiropoulos SN. An integrated approach to correction for off-resonance effects and subject movement in diffusion MR imaging. Neuroimage. 2016 Jan 15;125:1063–1078.

18. Smith SM, Jenkinson M, Woolrich MW, Beckmann CF, Behrens TEJ et al., Advances in functional and structural MR image analysis and implementation as FSL, Neuroimage 23 (2004) 208–219.

19. Grussu F, Schneider T, Tur C, Yates RL et al, Neurite dispersion: a new marker of multiple sclerosis spinal cord pathology?, Annals of Clinical and Translational Neurology (2017): vol. 4(9), pages 663–679.

20. Grussu F, Schneider T, Yates RL, Zhang H, Gandini Wheeler-Kingshott CAM, DeLuca GC and Alexander DC, A framework for optimal whole-sample histological quantification of neurite orientation dispersion in the human spinal cord, Journal of Neuroscience Methods (2016): vol. 273, pages 20–32.

21. Wang H, Lenglet C, Akkin T. Structure tensor analysis of serial optical coherence scanner images for mapping fiber orientations and tractography in the brain. J Biomed Opt. 2015 Mar; 20 (3):

22. Wang XT, Yu K, Wu SX, Gu JJ, Liu YH, Dong C, Qiao Y, Loy CC. ESRGAN: Enhanced Super-Resolution Generative Adversarial Networks. Computer Vision - Eccv 2018 Workshops, Pt V. 2019;11133:63–79.

23. Simonyan, Karen, and Andrew Zisserman. “Very deep convolutional networks for large-scale image recognition.” arXiv preprint 1409.1556 (2014).

24. Kingma, Diederik P., and Jimmy Ba. “Adam: A method for stochastic optimization.” arXiv preprint 1412.6980 (2014).

25. Li HY, Yang YF, Chang M, Chen SQ, Feng HJ, Xu ZH, Li Q, Chen YT. SRDiff: Single image super-resolution with diffusion probabilistic models. Neurocomputing. 2022;479:47–59.

